# Colouring coral larvae for tracking dispersal

**DOI:** 10.1101/2022.04.04.485987

**Authors:** Christopher Doropoulos, George Roff

## Abstract

1. Ecologists have long sought to understand larval dispersal characteristics of benthic marine invertebrates. Direct quantification of coral larvae dispersal has proven challenging, in part due to their complex life-history, minute size, and widespread dispersal at the scale of kilometres. Instead, indirect methods such as particle modelling, chemical signatures, and genetic correlation are often used in dispersal studies.
2. Here, we develop a direct method of quantifying larval dispersal by applying vital stains to coral larvae, allowing differentiation and direct tracking of millions of larvae from the pelagic dispersal stage through to the sedentary stages of attachment and metamorphosis on coral reefs.
3. Neutral red and Nile blue stains were extremely effective at staining coral larvae, while alizarin red and calcein blue showed no visible results. Differences in toxicity to vital stains was noted among species, with *Acropora* spp. exhibiting decreased larval survival and settlement, while Merulinidae spp. were unaffected. By experimenting with different incubation times and concentrations, our results indicate that neutral red can be effectively applied for short periods (<20 minutes) at low concentrations (1-100 mg l^-1^), whereas Nile blue requires longer stain times (>60 minutes) at higher concentrations (100-1000 mg l^-1^).
4. The strong colour of both neutral red and Nile blue stains was retained by newly settled larvae in lab settings upwards of five days following settlement, providing a direct method of differentiating between newly settled larvae on reefs. Field-validation of Nile blue applied to coral larvae from wild-captured coral slicks demonstrates the efficacy of staining across a diverse range of coral taxa.
5. Vital staining provides a simple, rapid (<60 mins), and low cost (<AUD$0.00001 per larva) method of colouring coral larvae that allows for direct tracking of dispersal and recruitment in studies of reef connectivity and restoration.

## 1. INTRODUCTION

Tracking and characterising the dispersal of benthic marine invertebrates is an age-old interest of marine ecologists (Thorson 1950). Benthic marine invertebrates have bipartite life-histories whereby mature sedentary adults release larval propagules into the plankton for a dispersive phase that can range from hundreds of metres to hundreds of kilometres, before returning to the benthos and undergoing metamorphosis into a new individual (Figure 1) (Cowen and Sponaugle 2009, Álvarez-Noriega et al. 2020). Reproduction often involves the release of eggs and sperm into the water column during mass spawning events where cross-fertilisation and larval development occurs, with the developing propagules typically <1 mm in size (Vanderklift et al. 2020). Due to this complex life-history, the minute size of propagules, and the extreme range of dispersal distances, characterising the origins and dispersal of benthic marine invertebrates continues to be a major challenge in marine ecology.

**Figure 1.**
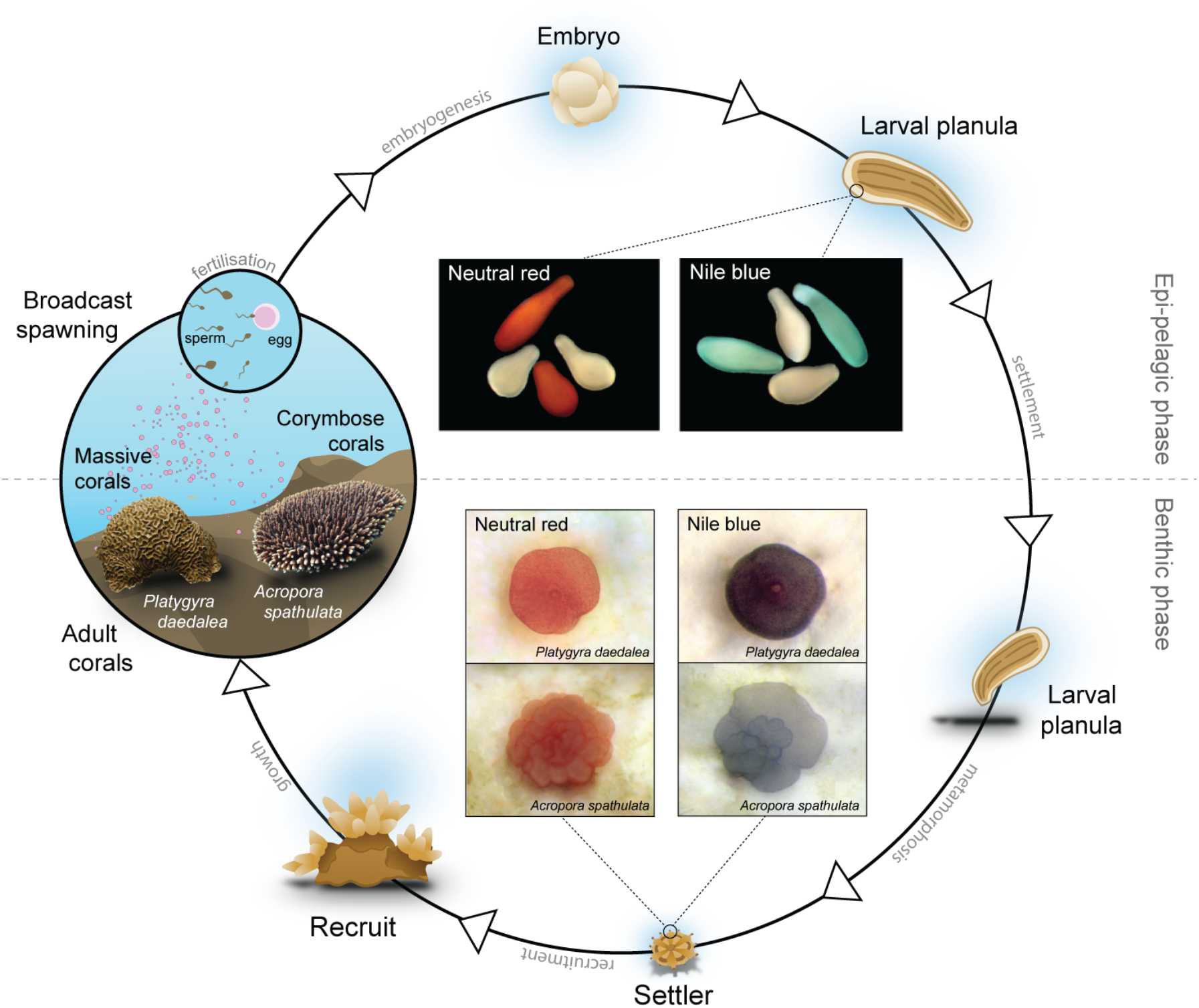
Life-history strategy of broadcast spawning corals, from the epi-pelagic release phase of sperm and eggs in mass spawning to the benthic stage of larval settlement and recruitment into the population. Inset photographs are representative images of neutral red and Nile blue stains on larvae (coloured vs control unstained larvae in white) and recently settled life-history phases.

Recent advances in physical modelling, simulated particle tracking and graph theory (Treml et al. 2007), elemental fingerprinting and chemical tracing (Becker et al. 2007), and genetic markers and population structure (Weersing and Toonen 2009), have been and continue to be at the forefront of characterising larval dispersal in recent years (Levin 2006). Such techniques operate across a broad range of spatial and temporal scales, but typically overlook finer scale processes and larval ecology (Metaxas and Saunders 2009) that ideally require direct quantification at the scale of individual larvae. One approach to overcome this limitation is larval staining, an approach that has been conducted since the early 1940’s (reviewed in Levin 1990). Larval staining offers a technique that can directly mark and recapture larvae for immediate and localised assessments, and incorporates larval behavioural ecology as well as local dispersal characteristics.

Like many benthic invertebrates, most corals (~84%) reproduce by broadcast spawning (Baird et al. 2009), whereby eggs and sperm are released into the water column where fertilisation, larval development and dispersal occurs (Babcock and Heyward 1986). Due to the ecological and economic importance of corals, coupled with the impacts of local stressors and climate change (Hughes et al 2017), research efforts have shifted towards the scaling of larval restoration (Doropoulos et al. 2019, Harrison et al. 2021, Vardi et al. 2021), with larval connectivity and conservation planning (Hock et al. 2017, Doropoulos and Babcock 2018, Mumby et al. 2021) at the forefront of research for management strategies to maintain the long-term resilience of coral reefs. This research has been hindered by the lack of direct methods to quantify larval connectivity and source-sink dynamics, with studies instead relying on indirect methods such as particle dispersal models and genetic correlation that operate at broader spatial and temporal scales.

Here, we develop a direct method of quantifying larval dispersal by applying vital stains to coral larvae that would allow for detection during i) the larval dispersal phase in the water column, or ii) during the sedentary phase following dispersal and settlement. The approach allows for empirical assessment of dispersal and connectivity potential, larval mortality rates, and rates of larval metamorphosis onto reefs. We first applied a broad array of stains at different concentrations and incubation times in an orthogonal approach using two common broadcast spawning corals in laboratory settings to understand general suitability, toxicity, and latent effects. In a second experiment, we took the two most effective stains and further refined incubation times and concentrations with an additional four species of common broadcast spawning corals. Finally, we applied the most effective and safest staining method on wild coral larvae in a field experiment to demonstrate the efficacy of vital stains as a method for colouring coral larvae in dispersal studies. Overall, the approach is an effective, rapid (<60 mins), and low cost (<$0.00001 per larvae) technique to directly track coral larvae and newly settled recruits in field studies.

## 2. MATERIALS AND METHODS

### 2.1 Vital stains

The following stains were used to test staining of coral larvae: Nile blue (C_20_H_20_ClN_3_O, CAS number: 3625-57-8) following the protocol of Allen et al. (2006); neutral red (C_15_H_17_N_4_, CAS number: 553-24-2) following the protocol of Loosanoff and Davis (1947); alizarin red (C_14_H_8_O_4_, CAS number: 72-48-0) following the protocol of Manzi and Donnelly (1971); and, calcein blue (C_15_H_15_NO_7_, CAS number: 54375-47-2) following the protocol of Tambutté et al. (2012).

### 2.2 Initial staining experiment

To establish the potential for larval staining, an initial factorial experiment was conducted using four stains at differing concentrations and incubation times. These experiments were conducted on two common species of coral, *Acropora spathulata* (family: Acroporidae, corymbose growth form) and coral *Platygyra daedalea* (family: Merulinidae, massive growth form). Both species were collected from the reef flat at Heron Island (southern Great Barrier Reef under permit number G19/42916.1) and maintained in 50 l aquaria prior to spawning. *A. spathulata* spawned on the 17^th^ November 2019 (21:45 - 00:40), and *P. daedalea* spawned on the 19^th^ of November 2019 (18:30 – 18:40). In both species, larval staining was conducted five days after spawning when larvae were developed (Figure 1).

*A. spathulata* larvae were placed with 10 mL stain in individual scintillation vials for 1, 6, 12, and 24 hr incubation periods at three different stain concentrations (1, 10, 100 mg l^-1^). For each of the vital stains, three replicates were conducted per each treatment (incubation time * stain concentration), with 20 larvae assigned to each replicate (36 treatments, 720 larvae total). Across all stains, this resulted in 144 treatments, totalling 2880 larvae. Staining was conducted in independent glass scintillation vials (10mL total volume). Larvae were added to each treatment so that the end point of the staining was the same across all treatments (i.e., larvae were added to the 1hr treatment at hr 23, 6hr treatment at hr 18, 12 hr treatment at hr 12), at which point larvae were 6 days old. The intensity of staining for each replicate was scored ordinally by a single observer (CD) into four categories: 1) no stain, 2) light staining, 3) medium staining, 4) strong staining. The proportion of alive larvae was counted under a dissecting microscope to determine the effects of staining on larval survival. To determine the effects of stain treatments on larval settlement, surviving larvae from each treatment were added to an individual conditioned settlement tile in 250 mL of filtered seawater. Water changes were conducted after 2 days (8 days after spawning), and larval settlement was scored 3 days after tiles were introduced (9 days after spawning).

*P. daedalea* larvae were placed with 20 mL stain in wells of cell-culture plates (Fiigure S1) for a single 36 hr incubation period at two different stain concentrations (10 and 100 mg l^-1^). For each of the four stains, a single replicate was conducted per each treatment (n = 150 larvae per each treatment, 4 treatments, 480 larvae total). Across all stains, this resulted in 16 treatments, and 1920 larvae total. Larval staining, scoring of stained larvae, and larval settlement followed the same protocol as for the *A. spathulata* experiment, with the exception of settlment, where larvae were settled on small chips (0.0625 cm^2^) of *Porlithon onkodes* crustose coralline algae (CCA) in cell-culture plates (100ml total volume).

To quantify significant differences between survival and settlement within each stain across different concentrations and incubation times for *A. spathulata*, we used a binomial Generalized Linear Model (GLM) for each stain, where incubation time and stain concentration were considered as fixed effects. Models were fit in R (v4.1.2) using the ‘glm’ function in the stats package (R Core Team 2021), and Tukey’s posthoc pairwise differences between treatments (incubation times and concentrations) were tested using the ‘glht’ function and visualised using the ‘cld’ functions in the multcomp package (Hothorn et al. 2008).

### 2.3 Refining staining experiment

To further refine the staining method, we conducted a follow-up experiment in November 2021 at the SeaSims aquaria facility (Australian Institute of Marine Science, Townsville, Australia). Based on the results of the initial experiments, two stains were discarded (alizarin red, calcein blue) and two were selected for further refining of incubation time and stain concentration (neutral red, Nile blue). To explore taxonomic differences in staining potential, four different coral species were used in the 2^nd^ experiment: *Acropora anthocersis* (spawning time: 22:30, 20^th^ October 2021), *Dipsastrea favus* (spawning time: 20:00, 23^rd^ October 2021), *Coelastrea aspera* (spawning time: 21:30, 24^th^ October 2021), and *Platygyra sinensis* (spawning time: 22:00, 24^th^ October 2021). These taxa are functionally distinct (tabular growth form: *A. anthocersis*, massive growth forms: *D. favus, P. sinensis, C. aspera*) and are phylogenetically distant (family: Acroporidae and family: Merulinidae). *A. anthocersis* neutral red staining was conducted at the following concentrations and incubation time treatments: 1mg l^-1^ for 15 mins, 10 mg l^-1^ for 10 & 30 mins, and for 100 mg l^-1^ for 5 and 10 mins, and a control (6 treatments, 180 larvae total). Nile blue staining was conducted at the following concentrations and incubation time treatments: 10, 100, and 500 mg l^-1^ for 60 mins, and for 500 mg and 1g l^-1^ for 120 mins (5 treatments, 150 larvae total). *G. aspera* neutral red staining was conducted at 10mg l^-1^ for 20 mins, and 100mg l^-1^ for 10 mins, and a control (3 treatments, 180 larvae total). Nile blue staining was conducted at 500 mg and 1g l^-1^ for 105 mins (2 treatments, 120 larvae total). *D. favus* neutral red staining was 650conducted at 10mg l^-1^ for 30 mins, and 100mg l^-1^ for 10 mins, and a control (3 treatments, 180 larvae total). Nile blue staining was conducted at 500 mg l^-1^ and 1 g l^-1^ for 120 mins (2 treatments, 120 larvae total). *P. sinensis* neutral red staining was conducted at 10mg l^-1^ for 20 mins, and 100mg l^-1^ for 10 mins (3 treatments, 180 larvae total). Nile blue staining was conducted at 500 mg l^-1^ and 1 g l^-1^ for 105 mins (2 treatments, 120 larvae total). Across all species, this resulted in 22 treatments and 1260 total larvae.

In each treatment, larvae were placed in 15 mL of stain solution in individual 6-well culture plates (Figure S1). Larval staining, scoring of stained larvae, and larval settlement followed the same protocol as for the first experiment, with small chips of *P. onkodes* CCA (0.0625 cm^2^) used to induce settlement. The intensity of stain staining for each replicate was scored ordinally by a single observer (CD) following the same protocol as the first experiment. To test for differences in larval survival and larval settlement among treatments, we used a binomial generalized linear model (GLM) using the ‘glm’ function in the stats package in R (version 4.1.2). Post-hoc differences between treatments (differing incubation times and concentrations) and controls were tested using the ‘glht’ function and visualised using the ‘cld’ function in the emmeans package (Lenth 2021).

### 2.4 Wild larval staining experiment

To quantify the efficacy of larval staining on natural wild collected larvae, we sampled multi-species coral slicks from the lagoon at Lizard Island (northern Great Barrier Reef, 14°41.045’S, 145°27.843’E) on the 4^th^ night after full moon (23^rd^ November 2021). Eggs were passively collected in the evening using a passive boom system and cultured in-situ in Lizard Island lagoon in a larval culture pool (5 x 5m). After 6 days, approximately 10,000 larvae were subsampled from the culture pool (estimated 1.5 million total) for larval staining. Larvae were stained with Nile Blue (Figure S1) to contrast the natural larval colours in a concentration of 1000 mg l^-1^ for 60 minutes. Stained and natural larvae were visually assessed under light microscopy.

## 3. RESULTS

### 3.1 Initial staining procedures

Larval staining of *Acropora spathulata* and *Platygyra daedalea* was highly effective using neutral red and Nile blue. These vital stains made the larvae and newly settled corals easily distinguishable compared to their natural colour (Figure 1). However, stain concentration and incubation times had toxic effects, which differed among stains and corals.

For *A. spathulata*, larval survival differed among stain, concentration, and incubation time (χ^2^ = 238.75, df=18, p < 0.001), with survival declining with increasing incubation time across all treatments (Figure 2). While the effects of the neutral red were most obvious at higher concentrations (10 & 100 mg l^-1^), stronger staining led to significantly lower larval survival at higher concentrations even at short incubation periods (Figure 2), with near complete larval mortality observed after 24hrs of incubation at higher concentrations. At the lowest concentration (1 mg l^-1^), neutral red achieved light staining after 6 hours, and medium levels of staining after 12 hours of incubation with >60% larval survival. Nile blue achieved the most consistent staining effect of all stains, with light stains observed at the lowest concentration (1 mg l^-1^) after a single hour of incubation (Figure 2). In contrast to the high mortality observed in the neutral red staining, survival was higher in Nile blue across all concentrations (Figure 2) and did not differ from controls at the 12hr time point (Figure S1). No significant differences were observed among low, medium, or high concentrations, and larval survival decreased with incubation time (Figure 2). Alizarin red and calcein blue failed to have any measurable effects on larvae regardless of concentration and incubation time. Larval survival did not differ from controls at the 12hr time point (Figure S2), and survival decreased with increasing incubation time in treatments (Figure 2).

**Figure 2.**
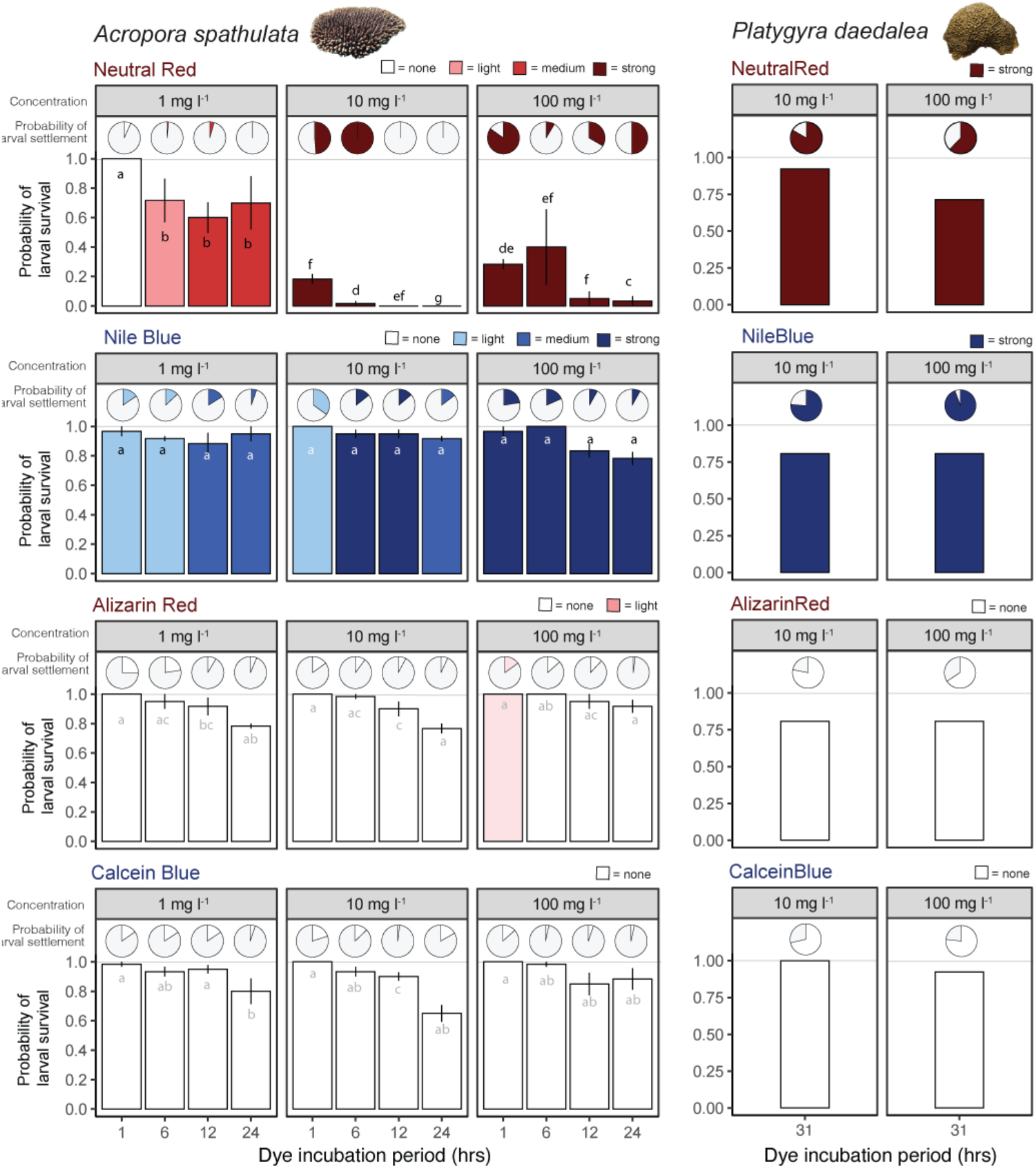
Probability of larval survival and settlement for two species of coral exposed to four stains at different concentration levels and incubation times. Colours indicate the strength of the larval stain at each stage (larval stage and settled larvae, inset key indicates none, light, medium or strong staining). Letters in bars indicate pairwise differences in probability of larval survival for *A. spathulata*. When a p value exceeds α = 0.05, then two means have at least one letter in common.

Larval settlement for *A. spathulata* differed significantly between differed among stain, concentration, and incubation time (χ^2^ = 227.49, df=18, p < 0.001), and was consistently low across all treatments (Figure 2). Of the initial larvae, most treatments resulting in <20% settlement, and more than half exhibiting less than 10% settlement (Figure 2). No significant differences were observed in the probability of settlement between control and stained larvae with the exception of neutral red, where settlement was significantly lower than the controls due to lower initial survival rates (Figure S2).

Larval survival of *P. daedalea* was considerably higher than *A. spathulata* (>75% survival, Figure 2), even with longer incubation times (31hrs vs 24hrs, Figure 2). Similar to *A. spathulata*, neutral red and Nile blue resulted in strong staining, at intermediate and high concentrations (10 & 100 mg l^-1^), while no staining effects of alizarin red and calcein blue were observed (Figure 2). In contrast to *A. spathulata*, larval settlement of *P. daedalea* was considerably higher, with >70% settlement rates across all stains and concentrations. Trials of mixed red and blue *P. daedalea* larvae placed with preconditioned settlement tiles showed that larval sources can be distinguished by utilising their colouration (Figure S3).

### 3.2 Refined staining procedures

The follow-up experiment confirmed the strong potential for neutral red and Nile blue stains across a range of functionally and phylogenetically distinct coral species (Figure S4). Under the refined staining procedures, larval survival was consistently high (>80%), with no significant difference among species and treatments (p > 0.05), nor to the controls (Figure 3).

**Figure 3.**
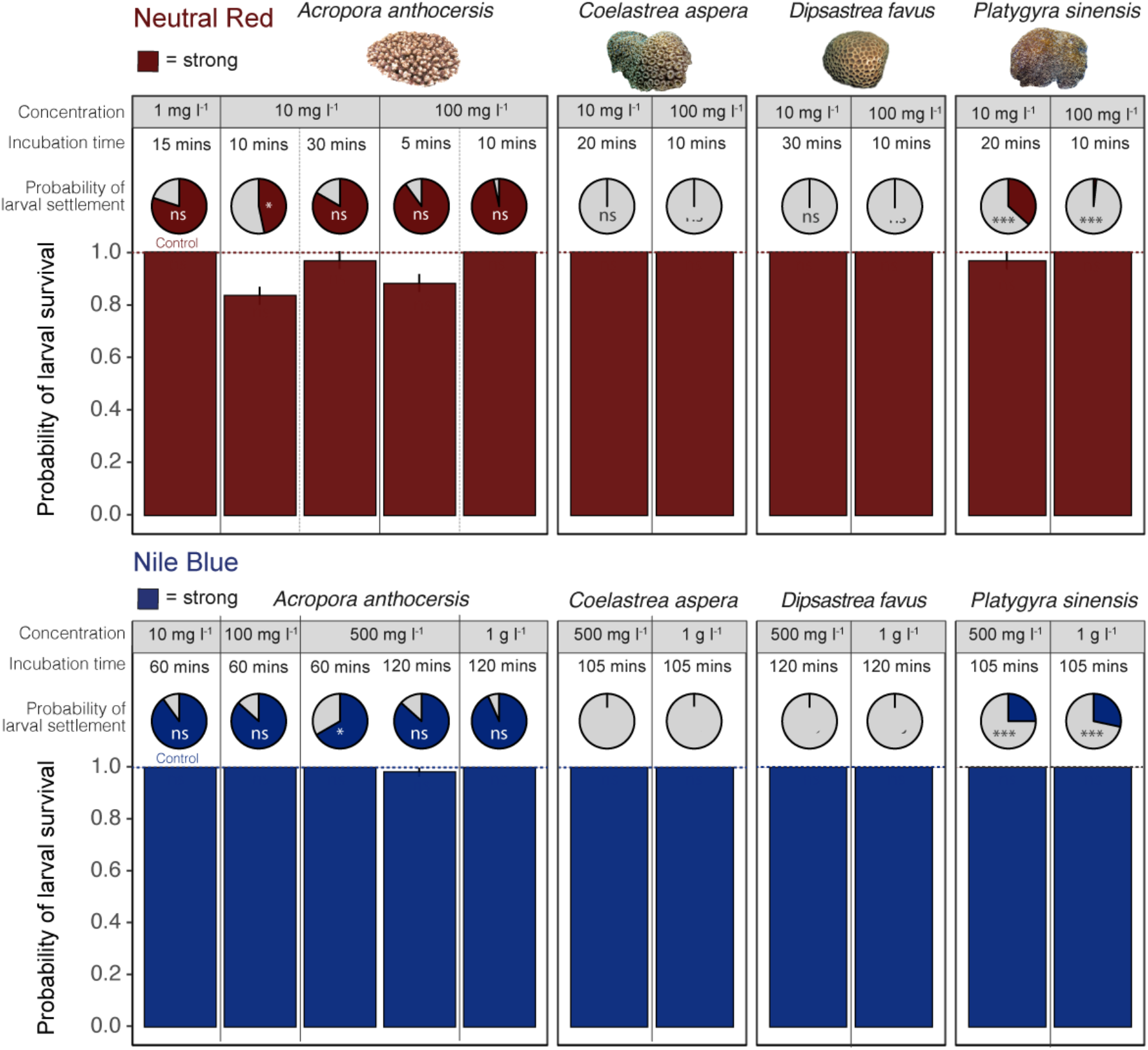
Probability of larval survival and settlement for four species of coral exposed to neutral red and Nile blue stains at different concentration levels and incubation times. Inset notation for probability of settlement indicates significant differences from control within each species at an α of 0.05 (^ns^ = no significant difference, * = p < 0.05, ** p < = 0.01, *** = p <0.001).

Larval settlement varied among species and treatments (χ^2^ = 46.2, df=6, p < 0.0001). For *Acropora anthocercis*, larval settlement was high (>75%) across all treatments and did not differ significantly from controls (p > 0.05), with an exception of a single treatment (500 mg l^-1^ for 60 mins) where settlement was reduced compared to the control (Figure 3). Larvae of *Goniastrea aspera* and *Dipsastraea favus* did not settle for the duration of the experiment in either Nile blue or neutral red. However, these results did not differ significantly from controls (p > 0.05) where no settlement was observed; suggesting the larvae were not competent and/or unresponsive to the CCA cue. Larval settlement of *Platygyra sinensis* in the neutral red stain was significantly higher in one treatment (10 mg l^-1^ for 20 mins), yet significantly lower in the other treatment (100 mg l^-1^ for 10 mins), indicating that higher stain concentrations may reduce larval settlement. Larval settlement of *P. sinensis* in the Nile blue stain was significantly higher in both treatments than in the control.

### 3.3 Staining wild coral larvae

Visual assessment of wild coral larvae under light microscopy confirmed the presence of a diverse multi-species larvae assemblage (size range: 150 - 650μm), ranging from cream - pink - red in colouration. Nile blue staining of these wild-captured larvae was highly effective, with >98% of larvae showing discernible staining effects (Figure 4). Following larval release onto a lagoonal patch reef, settlement of stained larvae was detected on settlement tiles. While the smaller cream-coloured larval species were effectively stained entirely blue in colour, the larger less common “bright-red” larval species appear to be stained blue only in the outer ectodermal layers, with the underlying, red-pigmented endodermal layers remaining partially visible (Figure 4).

**Figure 4.**
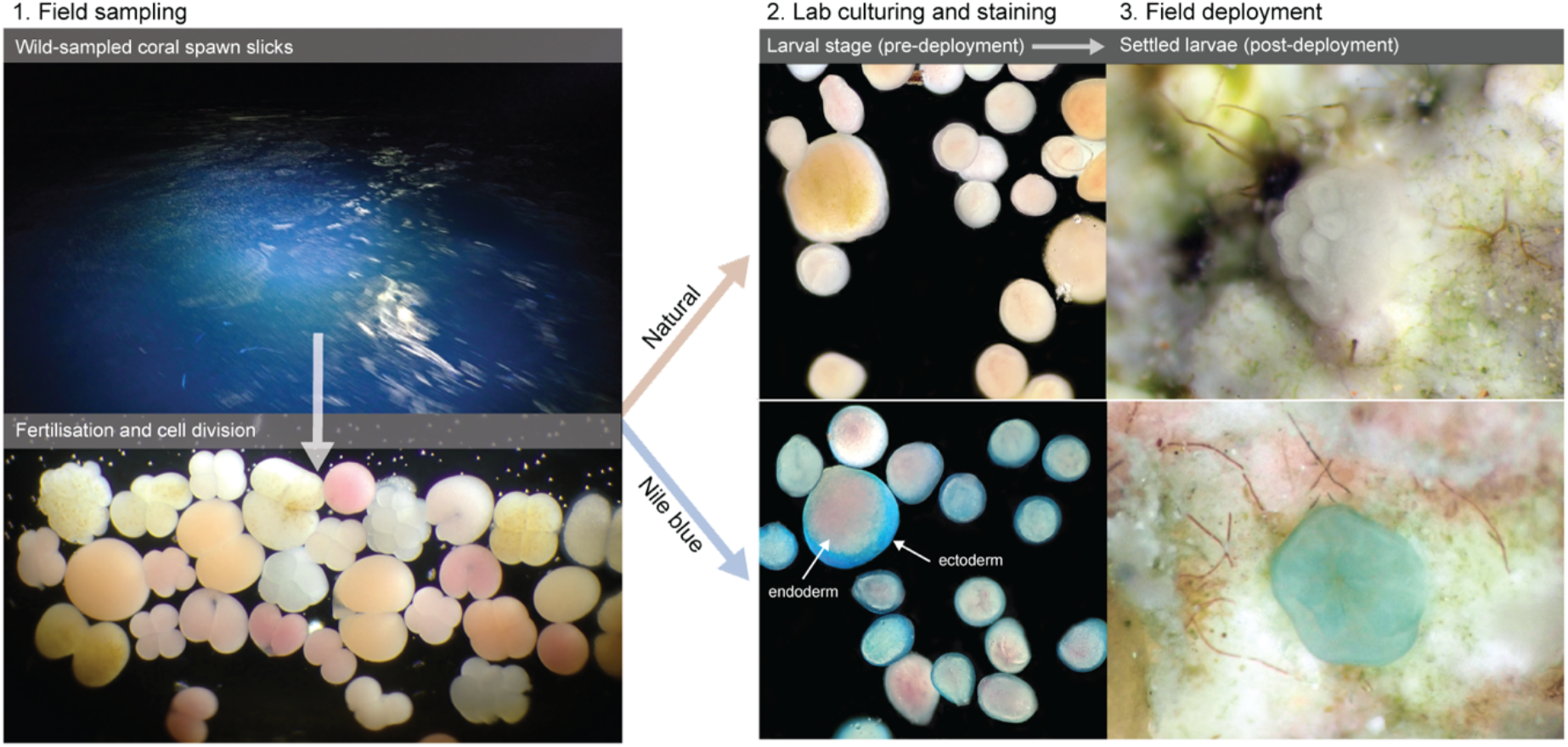
Field validation of larval staining: capture of wild-sampled coral spawn slicks showing high diversity of developing coral embryos, lab-based staining using Nile blue (1000 mg l^-1^ concentration for 60 minutes), and field deployment of competent stained and natural (unstained) larvae and detection on reef substrates.

## 4. DISCUSSION

We utilised a classic staining technique for benthic marine invertebrate larvae using histological stains and refined it for application to corals for tracking their dispersal as motile, epi-pelagic larvae and transition to sedentary, newly settled recruits on the benthos. Our first series of experiments worked across a range of stains, stain concentrations, and stain exposure times, to conduct initial investigations on effectiveness and critical thresholds utilising two common, functionally and phylogenetically distinct coral species. Staining potential was extremely high for two of the four stains – neutral red and Nile blue – yet differences in toxicity to the larvae at different stain concentrations and exposure times highlighted the need for procedural refinement. While we found that *Acropora* larvae mortality was more sensitive to neutral red, studies of other benthic marine invertebrate larvae have shown high sensitivity to Nile blue only (e.g., oysters, Manzi and Donnelly 1971) and not neutral red (Levin 1990). Our second series of experiments therefore optimised the stain concentrations and exposure times using neutral red and Nile blue, applying them at fewer concentrations and stain times, but working across an additional four coral taxa that are functionally and phylogenetically distinct. This second series of experiments was highly effective visually, mortality to larvae was extremely low, and settlement of stained and competent larvae high. Finally, we applied our staining technique to a culture of wild coral larvae following their capture after a synchronous mass spawning event and demonstrate its effectiveness in staining highly diverse communities of coral larvae as well as its potential to track the recruitment of newly settled coral larvae onto reefs. To our knowledge, we present the first application of staining to broadcast spawning coral larvae in laboratory and field studies to visually track their dispersal and recruitment on coral reefs.

The utilisation of our staining technique for tracking coral larvae dispersal and settlement has several advantages. Firstly, it is extremely simple, rapid, effective, and can be conducted in both laboratory and field settings. This allows for use in any kind of environment where access to specialised equipment may not be possible as is often the case in remote, tropical coral reefs. Secondly, the highly visual aspect of the stains makes it easy to see the coral larvae and settlers with the naked eye, even though the majority are only 300-900 μm in size (Babcock and Heyward 1986, Vanderklift et al. 2020). For detailed studies microscopy is required, but for rapid assessments inexpensive underwater cameras with macro capability can be used. Thirdly, histological stains are extremely accessible, being stocked by multiple major laboratory supply companies, and the concentrated powder forms allow for easy storage and transport to make bulk supplies on site. Both stains that we optimised are also extremely inexpensive: at optimal concentrations staining ~1 million larvae costs as little as ~AUD$10. As a comparison, other techniques such as molecular analysis for kinship (e.g. Cros et al. 2017) cost at a minimum ~$10 per individual and are far more time consuming in terms of sample processing (>20 hours). Finally, following our refinement of the stain concentration and exposure times, the staining assay causes minimal impacts on larval health or metamorphosis. Thus, even though the technique necessitates concentrating the larvae and staining for 5-120 minutes prior to release, it allows for the study of finer scale processes of larval ecology and behaviours such as rates of larval dispersal, larval mortality, larval supply-settlement relationships in field settings that other techniques typically overlook (Metaxas and Saunders 2009) or derive from laboratory experiments (e.g. Connolly and Baird 2010).

While effective in distinguishing individual larvae, staining approaches may have caveats that require further investigation. Importantly, brightly coloured particles in the water column may be an attractant for planktivorous fish, resulting in lower overall settlement rates. While some hard coral larvae are deeper red or greenish/blue in colouration, the majority are magenta-cream coloured (e.g., Figure 4 bottom left panel). Future studies could conduct feeding trials to quantify whether stained vs non-stained coral larvae alter preferential feeding by predators. Secondly, we have utilised the approach for short-term detection of coral settlers, up to 5 days following metamorphosis, but further studies are required to quantify the longevity of stained cells beyond this stage. Earlier studies applying neutral red and Nile blue on oysters and starfish have shown that stain retention can last anywhere from 6-70 days on larvae and up to 6 months following metamorphosis (Levin 1990). However, once newly settled corals calcify and uptake *Symbiodinium* in the first month following settlement (Abrego et al. 2009), the detectability of staining is likely to be reduced. Additionally, as the coral recruit begins to grow and asexually divide, the detectability of staining may dilute as the new polyps form. Future work could conduct longer-term assessments of detectability as newly settled corals develop to assess the durability of the approach for longer-term post-settlement studies that could go beyond larval dispersal and initial recruitment.

In summary, we provide a methodology for colouring coral larvae for tracking the dispersal and settlement of broadcast spawning corals. The approach can be applied to distinct lines of coral species from diverse functional and phylogenetic lineages, as well as diverse communities spawned from natural communities. From an experimental perspective, the ability to use different colour stains can allow differentiation between larval cohorts to examine fine scale behavioural responses and interactive processes. At larger scales, the approach may be useful to help validate simulation studies of local coral connectivity (Hock et al. 2017, Doropoulos and Babcock 2018, Mumby et al. 2021) and differentiate between targetted larval releases and background settlement during large-scale restoration programs (Doropoulos et al. 2019, Harrison et al. 2021, Vardi et al. 2021) with a view to optimising management strategies for conservation planning of coral reefs.

## ACKNOWLEDGEMENTS

The authors would like to acknowledge the Traditional Owners of the Great Barrier Reef, particularly the Gidarjil First Nations people, the Bindal, Wulgurukaba and Manbarra First Nations people, and the Ngurruumungu and Dingaal First Nations people, for permission to collect corals and bring them to laboratory facilities in their Sea Country. We thank Russell McCulloch from CSIRO Agriculture & Food for providing us with the stains; Pascal Craw, Jesper Elzinga, Lauren Hardiman, Damian Thomson and the Moving Corals team for field assistance; Andrea Severati, Muhammad Azmi Abdul Wahab and SeaSIM staff for logistical support; and Russ Babcock and Carly Randall for useful discussions. This research was funded by CSIRO Oceans & Atmosphere and the Reef Restoration and Adaptation Program (RRAP). RRAP is funded by the partnership between the Australian Government’s Reef Trust and the Great Barrier Reef Foundation

## AUTHOR CONTRIBUTIONS

CD conceived and designed the study; CD and GR performed the study; GR analysed the data; GR and CD co-led writing the manuscript.

## DATA AVAILIBILITY STATEMENT

Data available at https://data.csiro.au/ and R code available at https://github.com/marine-ecologist.

---

**Figure S1.**
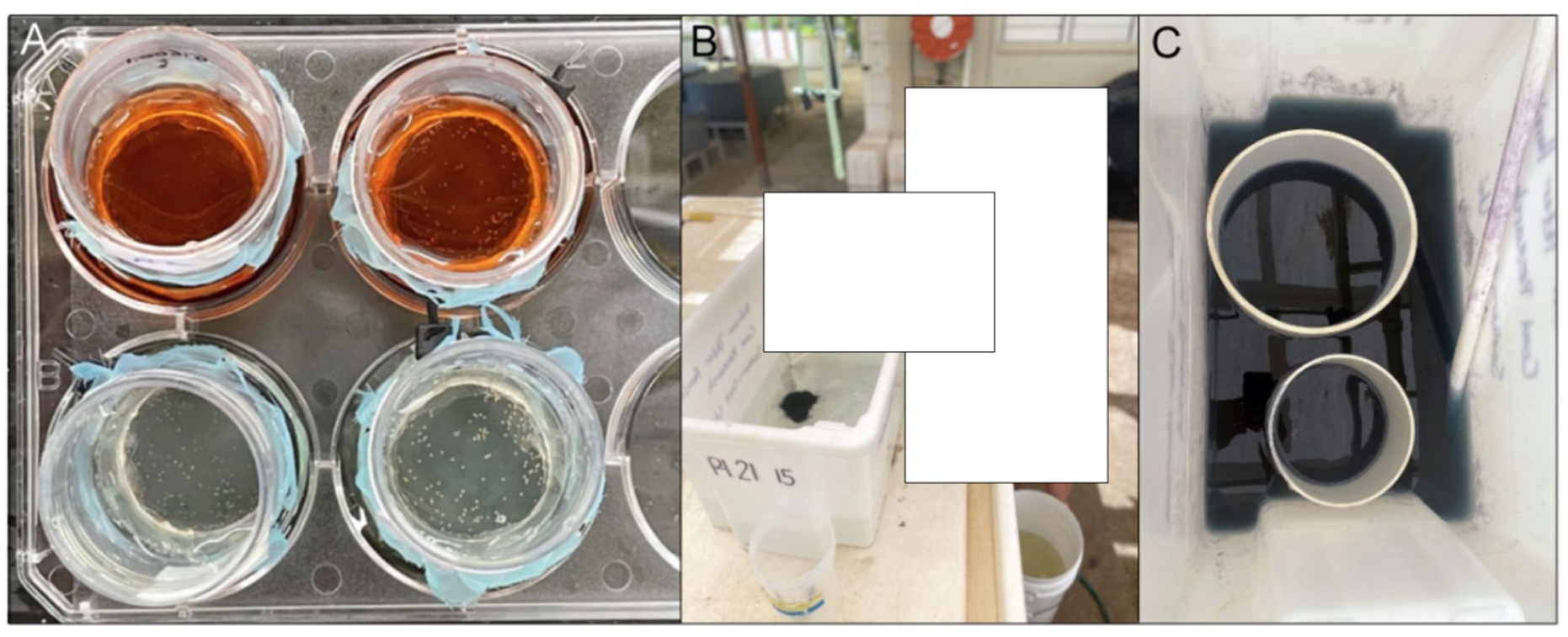
Procedural approaches to stain larvae at laboratory and field scales. a) Staining coral larvae in small separators that are nesting in varying concentrations of neutral red and Nile blue solutions in 6-well cell-culture plate wells for easy removal at different times and rinsing following removal. b) Mixing of Nile blue staining in seawater into which c) larvae are retained in the stain within large separators for easy removal and rinsing prior to deployment.

**Figure S2.**
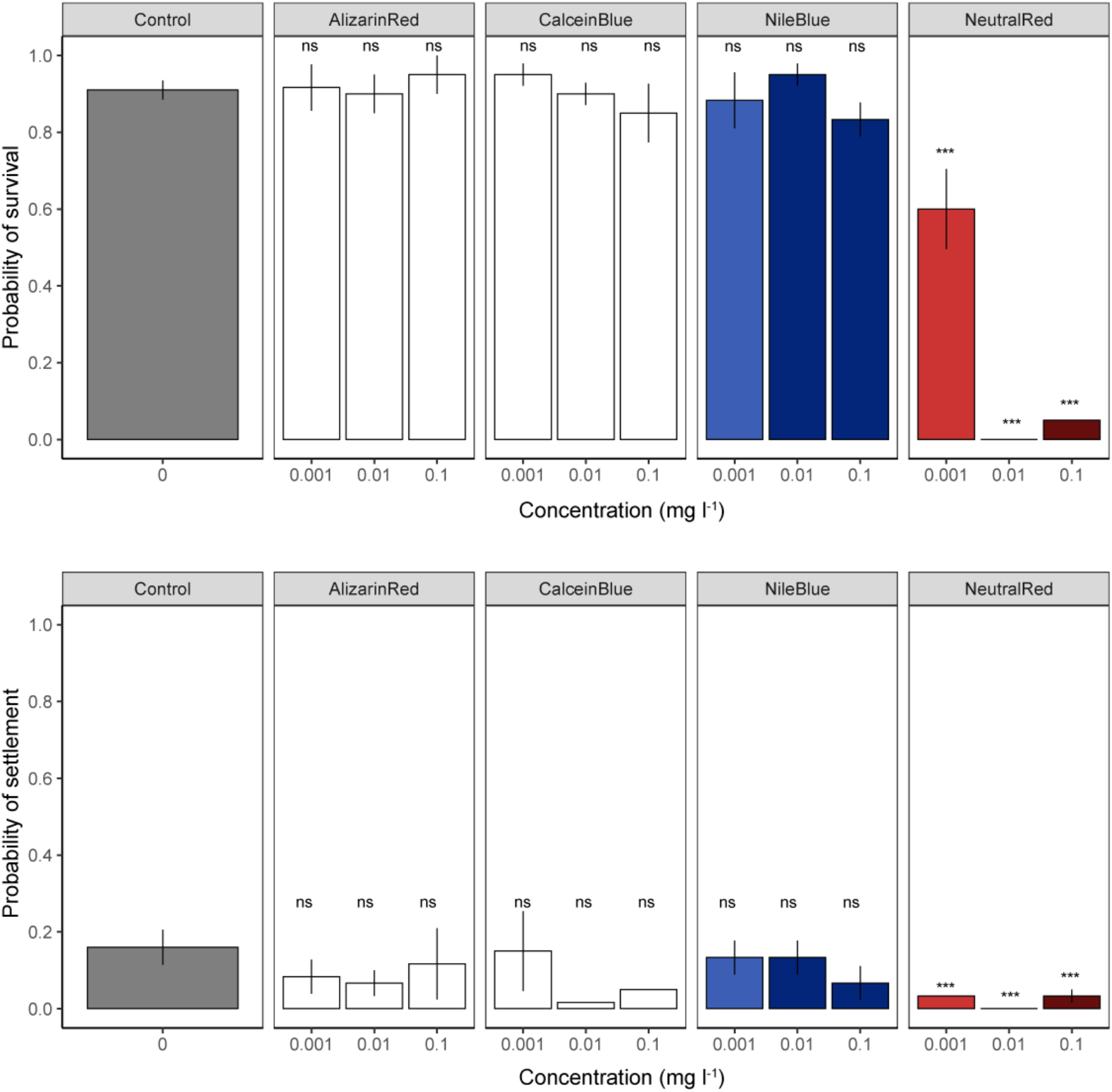
Probability of larval survival and larval settlement for *Acropora spathulata* exposed to four stains (neutral red, Nile blue, alizarin red, calcein blue) at different concentration levels after 12 hours of incubation and control (unstained) larvae. Colours indicate the strength of the larval stain (see Figure 2 for legend). Pairwise differences indicate significant differences from control (^ns^ = no significant difference, * = p < 0.05, ** p < = 0.01, *** = p <0.001).

**Figure S3.**
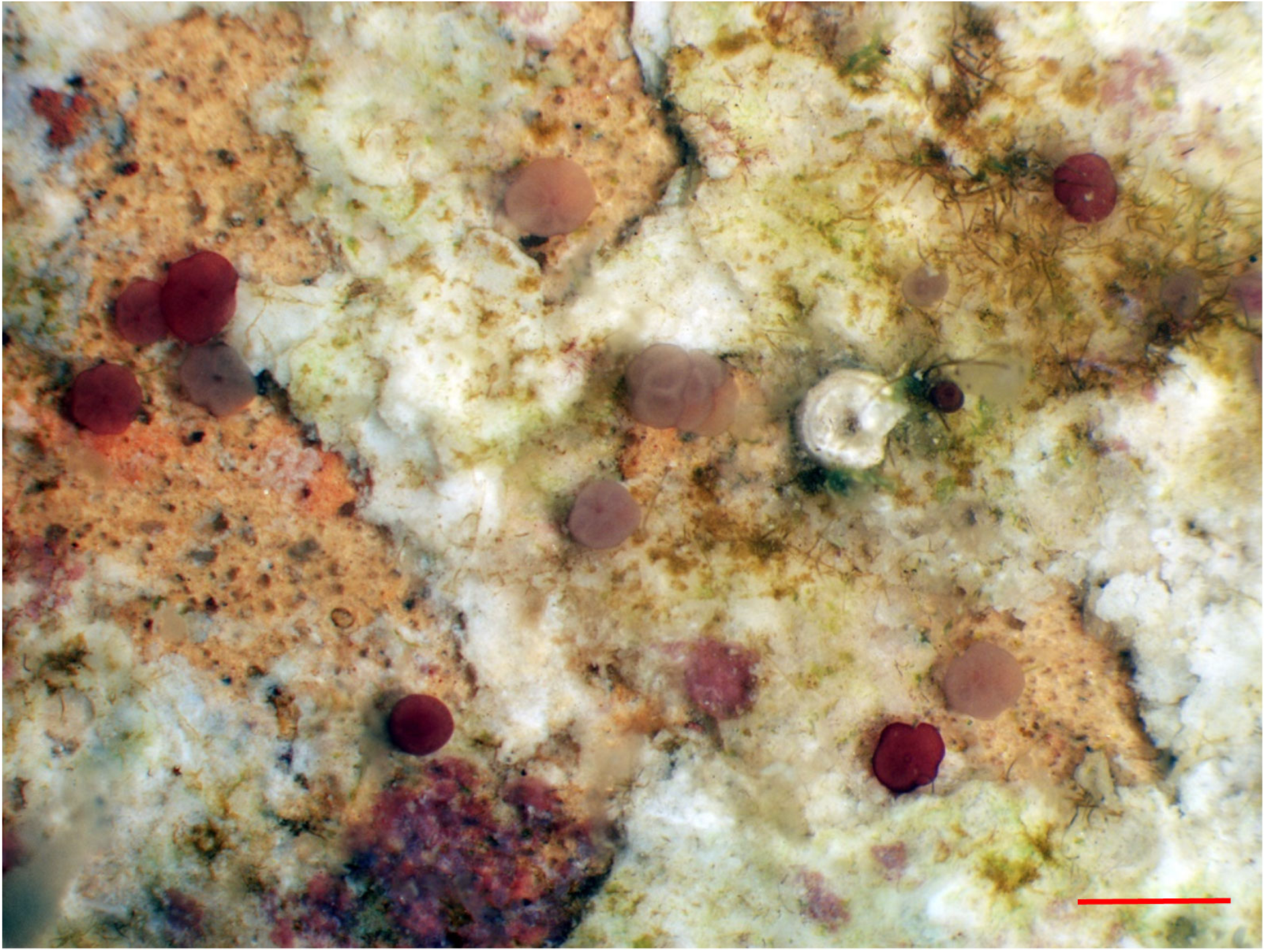
Example of a settlement tile with newly settled *Platygyra daedalea* 8 days after spawning following a mixed staining treatment of 50% neutral red stain, 50% Nile blue stain under a light microscope. Red scale bar = 1 mm

**Figure S4.**
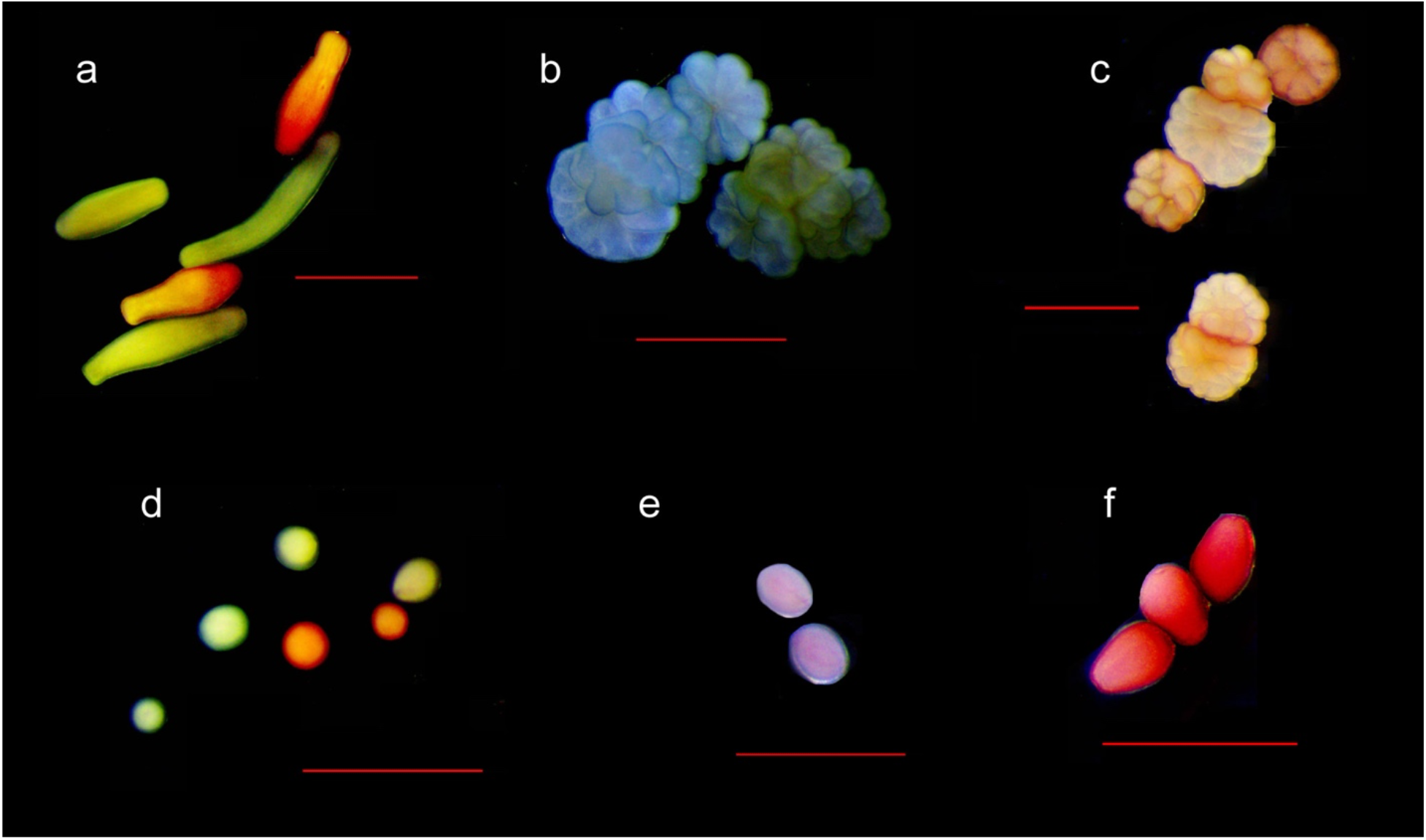
Additional examples of stained larvae and newly metamorphosed spat from common spawning coral species: a) *Acropora anthocercis* larvae (Nile blue, neutral red), b) *Acropora anthocercis* spat (Nile blue), c) *Acropora anthocercis* spat (neutral red), d) *Dipsastraea favus* (Nile blue, neutral red), d) *Platygyra sinensis* (Nile blue), e) *Platygyra sinensis* (neutral red). Red scale bars = 1 mm.

